# Microstimulation in a spiking neural network model of the midbrain superior colliculus

**DOI:** 10.1101/424127

**Authors:** Bahadir Kasap, A. John van Opstal

## Abstract

The midbrain superior colliculus (SC) generates a rapid saccadic eye movement to a sensory stimulus by recruiting a population of cells in its topographically organized motor map. Supra-threshold electrical microstimulation in the SC reveals that the site of stimulation produces a normometric saccade vector with little effect of the stimulation parameters. Moreover, electrically evoked saccades (E-saccades) have kinematic properties that strongly resemble natural, visual-evoked saccades (V-saccades). These findings support models in which the saccade vector is determined by a center-of-gravity computation of activated neurons, while its trajectory and kinematics arise from downstream feedback circuits in the brainstem. Recent single-unit recordings, however, have indicated that the SC population also specifies instantaneous kinematics. These results support an alternative model, in which the desired saccade trajectory, including its kinematics, follows from instantaneous summation of movement effects of all SC spike trains. But how to reconcile this model with microstimulation results? Although it is thought that microstimulation activates a large population of SC neurons, the mechanism through which it arises is unknown. We developed a spiking neural network model of the SC, in which microstimulation directly activates a relatively small set of neurons around the electrode tip, which subsequently sets up a large population response through lateral synaptic interactions. We show that through this mechanism the population drives an E-saccade with near-normal kinematics that are largely independent of the stimulation parameters. Only at very low stimulus intensities the network recruits a population with low firing rates, resulting in abnormally slow saccades.

**Author Summary:** The midbrain Superior Colliculus (SC) contains a topographically organized map for rapid goal-directed gaze shifts, in which the location of the active population determines size and direction of the eye-movement vector, and the neural firing rates specify the eye-movement kinematics. Electrical microstimulation in this map produces eye movements that correspond to the site of stimulation with normal kinematics. We here explain how intrinsic lateral interactions within the SC network of spiking neurons sets up the population activity profile in response to local microstimulation to explain these results.

## 1 Introduction

High-resolution foveal vision covers only 2% of the visual field. Thus, the visual system has to gather detailed information about the environment through rapid goal-directed eye movements, called saccades. Saccades reach peak eye velocities well over ~1000 deg/s in monkey, and last for only 40-100 ms, depending on their size. The stereotyped relationships between saccade amplitude and ~duration (described by a straight line) and ~peak eye velocity (a saturating function) are termed the ‘saccade main sequence’ (Bahill et al., 1975). The acceleration phase of saccades has a nearly constant duration for all amplitudes, leading to positively skewed velocity profiles (Van Opstal et al., 1987). In addition, the horizontal and vertical velocity profiles of oblique saccades are coupled, such that they are scaled versions of each other (through component stretching), and the resulting saccade trajectories are approximately straight (Van Gisbergen et al., 1985). These kinematic properties all imply that the saccadic system contains a nonlinearity in its control (Van Gisbergen et al., 1981, 1985; Smit et al., 1990). More recent theories hold that this nonlinearity reflects an optimization strategy for speed-accuracy trade-off, which copes with the spatial uncertainty in the retinal periphery, and internal noise in the sensorimotor pathways (Harris and Wolpert, 1998; Tanaka et al., 2006; Van Beers, 2008; Goossens and Van Opstal, 2012).

The neural circuitry responsible for saccade programming and execution extends from the cerebral cortex to the pons in the brainstem. The midbrain superior colliculus (SC) is the final common terminal for all cortical and subcortical inputs, and it has been hypothesized to specify the vectorial eye-displacement command for downstream oculomotor circuitry (Robinson, 1972; Scudder, 1988; Moschovakis, 1998). The SC contains an eye-centered topographic map of visuomotor space, in which the saccade amplitude is mapped logarithmically along its rostral-caudal anatomical axis (*u*, in mm) and saccade direction maps roughly linearly along the medial-lateral axis (*v*, in mm; Robinson, 1972). The afferent map (Eqn. 1a) and its efferent inverse (Eqn. 1b) can be described by (Ottes et al., 1986):

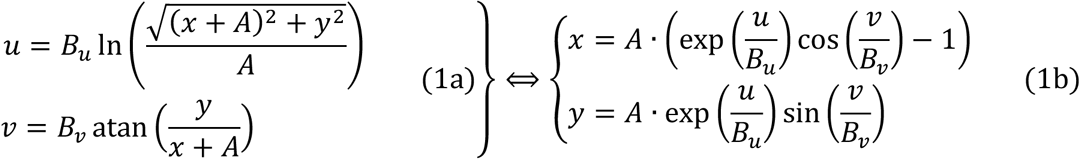

with parameters B_u_≈1.4 mm, B_v_≈1.8 mm/rad, and A≈3 deg. Each saccade is associated with a translation-invariant Gaussian-shaped population within this map, the center of which corresponds to the saccade vector, (*x,y*), and a width of ***σ***≈0.5 mm (Ottes et al., 1986; Van Opstal et al., 1990). It is generally assumed that each recruited neuron, *n*, in the population encodes a vectorial movement contribution to the saccade vector, which is determined by both its anatomical location within the motor map, (*u_n_*, ***v***_*n*_), and its activity, *F_n_*.

Precisely how individual cells contribute to the saccade is still debated in the literature. Two competing models have been proposed for decoding the SC population: weighted averaging of the cell vector contributions (Lee et al., 1988; Port and Wurtz, 2003; Walton et al., 2005; Eqn. 2a) vs. linear summation (Van Gisbergen et al., 1985; Goossens and Van Opstal, 2006, 2012; Eqn. 2b), respectively, which can be formally described as follows:

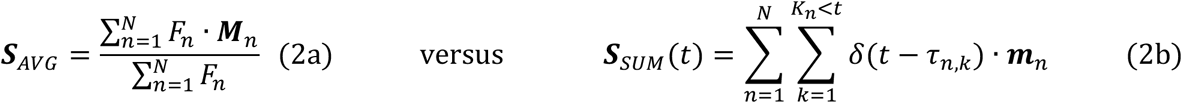

*N* is the number of active neurons in the population, *K_n_*<t the number of spikes in the burst of neuron *n* up to time *t*, *F_n_* its mean (or peak) firing rate, and ***M***_*n*_=*(x_n_,y_n_)* is the saccade vector in the motor map encoded at SC site *(u_n_,**v**_n_)* (Eqn. 1b).

**m**_n_ = ζ**M**_n_ is the small, fixed vectorial contribution of cell *n* in the direction of **M**_n_, for each of its spikes, with ζ a fixed, small scaling constant that depends on the adopted cell density in the map and the population size, and *δ(t*-***τ****k,n)* is the *k*’th spike of neuron *n*, fired at time τ_k,n_.

The vector-averaging scheme of Eqn. (2a) only specifies the amplitude and direction of the saccade vector, and thus puts the motor map of the SC outside the kinematic control loop of its trajectory. It assumes that the nonlinear saccade kinematics are generated by the operation of horizontal and vertical dynamic feedback circuits in the brainstem (Jürgens et al. 1981, Robinson 1975; Lee et al., 1988), or cerebellum (Lefèvre et al 1998, Quaia et al. 1999). Note also that vector averaging is a nonlinear operation because of the division by the total population activity.

In contrast, the linear dynamic ensemble-coding model of Eqn. (2b) encodes the full kinematics of the desired saccade trajectory at the level of the SC motor map through the temporal distribution of spikes by all cells in the population (Goossens and Van Opstal, 2006; 2012). As a result, the instantaneous firing rates of all neurons in the population, *f_n_(t)*, together encode the desired vectorial saccadic velocity profile:

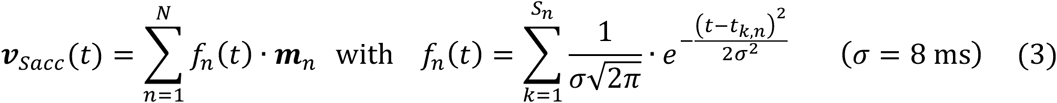

where *S_n_* is the number of spikes of cell *n*, with the spikes occurring at times *t_k,n_*.

Although the models of Eqn. 2a,b cannot both be right, each is supported by different lines of evidence. For example, electrical microstimulation produces fixed-vector (E-)saccades with normal main-sequence kinematics that are insensitive to a large range of stimulation parameters (Robinson, 1972; Van Opstal et al., 1990; Stanford et al., 1993; Katnani and Ghandi, 2012). If one supposes that electrical stimulation directly activates a large population of SC cells, and that the firing rates follow the (typically rectangular) stimulation profile, a vector-averaging scheme with downstream dynamic feedback circuitry readily explains why E-saccades are normal main-sequence, since the center of gravity of the population specifies the desired saccade vector only, regardless the firing rates.

In addition, reversible inactivation of a small part of the SC motor map produces particular deficits in the metrics of visually-evoked (V-)saccades that may not be readily explained by the linear summation model of Eqn. 2b (Lee et al., 1988). As the amplitude and direction of a V-saccade to the center of the lesioned site remain unaffected, saccades to locations around that site are directed away from the lesion. For example, V-saccades for sites rostral to the lesion undershoot the target, while V-saccades for sites caudal to the lesion will overshoot the target.

The simple vector-summation model of Eqn. 2b yields saccades that would always undershoot targets, as the lesioned population produces fewer output spikes than under normal control conditions. However, Goossens and Van Opstal (2006, 2012) observed that their estimate of the total number of spikes from the SC population, was remarkably constant, regardless saccade amplitude, direction, or speed. Yet, they also observed that many cells in the normal SC fire some post-saccadic spikes. They therefore assumed that saccades are actively terminated by a downstream mechanism, whenever the criterion of a fixed number of spikes, NTOT, is reached:

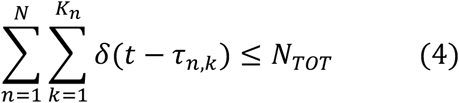

They demonstrated, by simulating the summation model of Eqn. 2b with actual recordings from ~150 cells, that by including the criterion of Eqn. 4 (which constitutes a cut-off nonlinearity in the model), the pattern of saccadic over- and undershoots to a focal SC lesion can be fully explained. In addition, the extended summation model of Eqns. 2b and 4 also accounts for weighted averaging of double-target stimulation in the motor map (Robinson, 1972; Van Opstal and Van Gisbergen, 1989; Van Opstal, 2016). Moreover, although the vector-averaging model (Eqn. 2a) correctly predicts the pattern of saccadic dysmetrias, it fails to explain the substantial slowing of the lesioned saccades (Lee et al., 1988). As this latter observation is also accounted for by Eqns. 2b and 4 (Goossens and Van Opstal, 2006), it further supports the hypothesis that the SC population encodes both the saccade-vector, and its kinematics.

Interestingly, electrical microstimulation experiments have also shown that at low current strengths, just above threshold, the evoked saccade vectors become smaller and slower than main sequence (Van Opstal et al., 1990; Katnani and Ghandi, 2012). These results do not follow from vector averaging but are readily predicted by dynamic summation (Eqns. 2b and 4).

However, if microstimulation would produce a large square-pulse population profile around the electrode tip (mimicking the profile of the imposed current pulses, as is typically assumed), the summation model would generate severely distorted saccade-velocity profiles, which are not observed in experiments. Yet, little is known about the actual activity profiles in the motor map evoked by electrical microstimulation, as simultaneous multi-electrode recordings during microstimulation are not available and would be obscured by the large stimulation artefacts (Histed et al., 2013).

Under microstimulation, two factors contribute to neuronal activation: (1) direct (feedforward) current stimulation of cell bodies and axons by the stimulation pulses of the electrode, and (2) synaptic activation through lateral (feedback) interactions among neurons in the motor map. How each of these factors contributes to the population activity in the SC is unknown. It is conceivable, however, that current strength falls off rapidly with distance from the electrode tip (at least by ~1/r^2^), and that hence a relatively small number of SC neurons would be directly stimulated by the electric field of the electrode.

Indeed, a recent two-photon imaging study, carried out in frontal eye fields (FEF), showed that microstimulation at physiological current strengths activates only a sparse set of neurons directly around the immediate vicinity of the stimulation site (Histed et al., 2009). These considerations therefore suggest that the major factor in explaining the effects of microstimulation in the SC motor map may be synaptic transmission through lateral excitatory-inhibitory connections among the cells. Such a functional organization in the SC is supported by anatomical studies (Behan and Kime, 1996; Olivier et al., 1998), by electrophysiological evidence (Munoz and Istvan, 1998; Phongphanphanee et al., 2011; 2014), and by pharmacological studies (Meredith and Ramoa, 1998).

We recently constructed a biologically plausible, yet simple, spiking neural network model for ocular gaze-shifts by the SC population to visual targets (Kasap and Van Opstal, 2017). This minimalistic (one-dimensional) model with lateral interactions can account for the experimentally observed firing properties of saccade-related cells in the gaze-motor map (Goossens and Van Opstal, 2006, 2012), by assuming an invariant input pattern from sources upstream from the motor map (e.g., FEF).

We here extended that simple spiking neural network model to account for microstimulation results over a wide range of stimulation parameters, and to generate appropriate saccadic command signals across the two-dimensional oculomotor range.

## 2. Methods

### 2.1 Log-polar afferent mapping

The afferent mapping function (Eqn. 1a) translates a target point in visual space to the anatomical position of the center of the corresponding Gaussian-shaped population in the SC motor map. It follows a log-polar projection of retinal coordinates onto Cartesian collicular coordinates (Eqn. 1a; Ottes et al., 1986). To allow for a simple 2D matrix representation of the map in our network model, we simplified the afferent motor map to the complex logarithm:

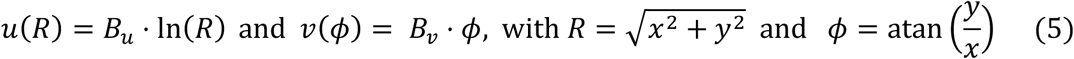

with *B_u_* = 1 mm and *B_v_* = 1 mm/rad (isotropic map). Thus, the contribution, ***m***, of a single spike at site (*u,v*) to the eye movement is computed from the efferent mapping function as:

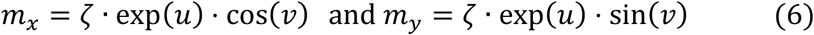

We thus constructed a spiking neural network model as a rectangular grid of 201 × 201 neurons. The network represents the gaze motor-map with 0 < *u* < 5 mm (i.e., up to amplitudes of 148 deg), and 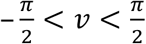 mm. The network generates saccadic motor commands of different directions and amplitudes into the contralateral visual hemispace through a spatial-temporal population activity profile. The location of the population in the motor map determines the direction and amplitude of the saccade target, whereas the temporal activity profile encodes the eye-movement kinematics, through Eqn. 2b. As described below, and in our previous study (Kasap and Van Opstal, 2017), the eye-movement main-sequence kinematics result from location-dependent biophysical properties of the neurons within the map, together with their lateral interconnections.

### 2.2 The AdEx neuron model

We investigated the dynamics of the network model numerically through simulations developed in C++/CUDA (Nickolls et al., 2008). The motor map is represented as a rectangular grid of neurons with a Mexican hat-type pattern of lateral interactions. The neural activities were simulated by custom code utilizing dynamic parallelism to accelerate spike propagation on a GPU (Kasap and Van Opstal, 2018). The code was developed and tested on a Tesla K40 with CUDA Toolkit 7.0, Linux Ubuntu 16.04 LTS (repository under https://bitbucket.org/bkasap/sc_microstimulation). Simulations ran with a time resolution of 0.01 ms. Brute-force search and genetic algorithms, described below, were used for parameter identification and network tuning since there exists no analytical solution for the system.

The neurons in the network were described by the adaptive exponential integrate-and-fire (AdEx) neuron model (Brette and Gerstner, 2005), which accommodates for a variety of bursting dynamics with a minimum set of free parameters. The AdEx model is a conductance-based integrate-and-fire model with an exponential membrane potential dependence. It reduces Hodgkin-Huxley’s model to only two state variables: the membrane potential, *V*, and an adaptation current, *q*. The temporal dynamics of the system are given by the following differential equations for neuron *n*:

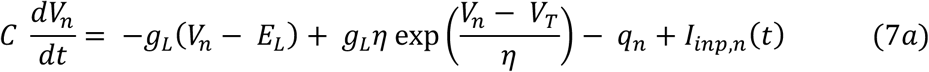

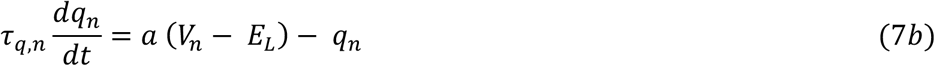

where *C* is the membrane capacitance, *g_L_* is the leak conductance, *E_L_* is the leak reversal potential, *η* is a slope factor, *V_T_* is the neural spiking threshold, *τ_q_* is the adaptation time constant, *a* is the sub-threshold adaptation constant, and *I_inp,n_* is the total synaptic input current. In our previous paper (Kasap and Van Opstal, 2017) the input-layer of Frontal Eye Field (FEF) neurons had identical biophysical properties, and only received a fixed external input current, *I_inp,n_* = *I_ext_*. In the present simulations, we did not include a FEF input layer, as the electrical stimulation was applied within the SC motor map as an external current.

Two parameters specify the biophysical properties of the SC neurons: the adaptation time constant, *τ_q,n_* (which is assumed to be location dependent), and the synaptic input current, *I_inp,n_* = *I_syn,n_* + (where *I_syn,n_* is a location- and activity-dependent synaptic current, and *I_E_* is the applied microstimulation current). Both variables change systematically with the spatial location of the cells within the network (rostral to causal). The remaining parameters, *C*, *g_L_*, *E_L_*, *η*, *V_T_* and *a*, were tuned such that the cells showed neural bursting behavior (see Table 1 for the list and values of all parameters used in the simulations, and Fig. 1 for some example responses).

**Figure 1:**
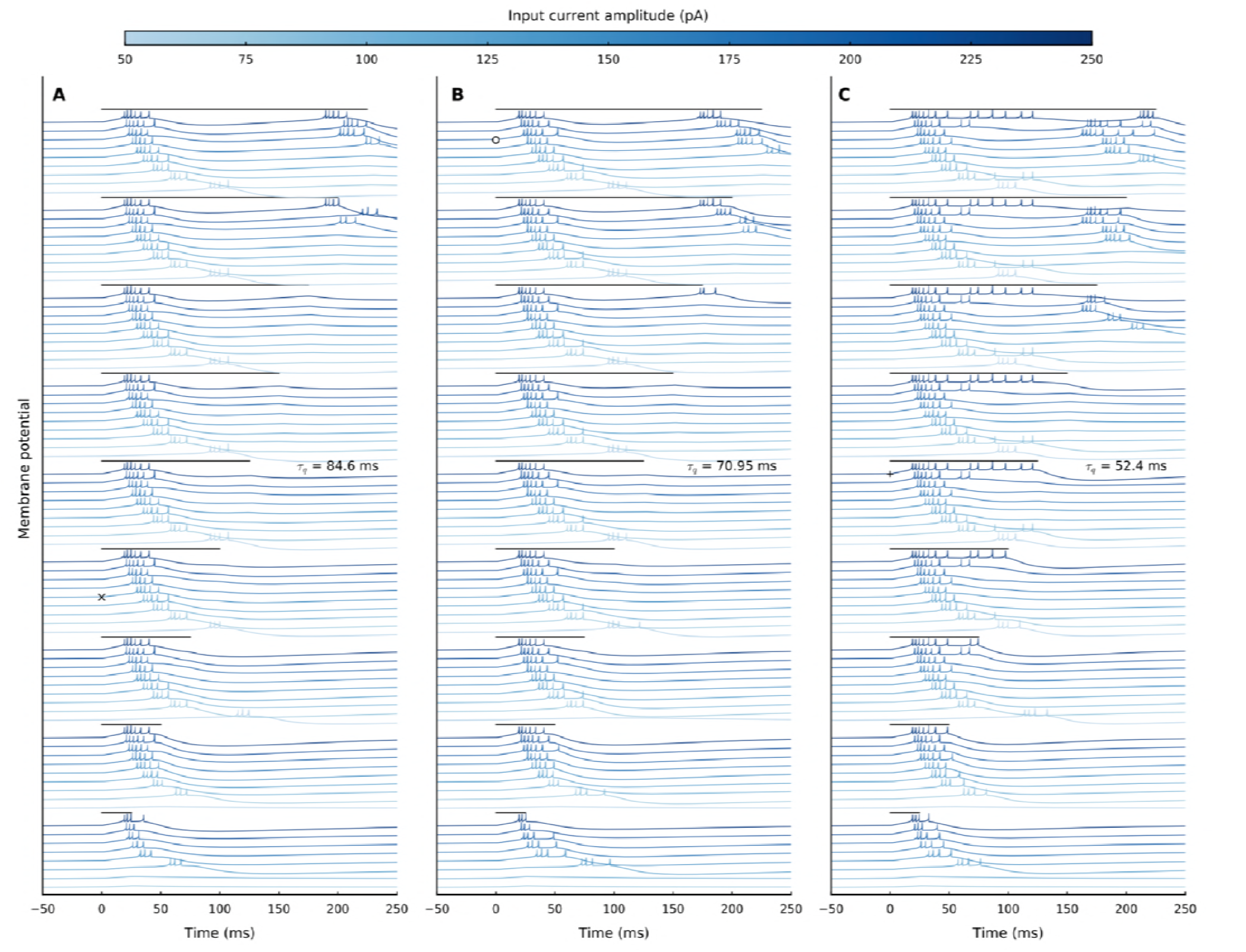
Responses of three SC model neurons to different microstimulation parameters. The three neurons differed in their adaptation time constants (**A**: τ_q_ = 84.6 ms, **B:** τ_q_ = 70.95 ms, and **C**: τ_q_ = 52.4 ms). Each row shows the membrane potentials, V(t), for the same electrical stimulus, at a particular intensity (see color code for the different lines, top), and delivered at a particular stimulus duration, DS. Note the clear differences in neuronal membrane responses. Stimulus timings and durations are indicated above the traces by black lines, ranging from DS=25 ms (bottom) to DS=225 ms (top). Symbols x, o, and +: selected responses, further analyzed in Fig. 2.

**Table 1.**
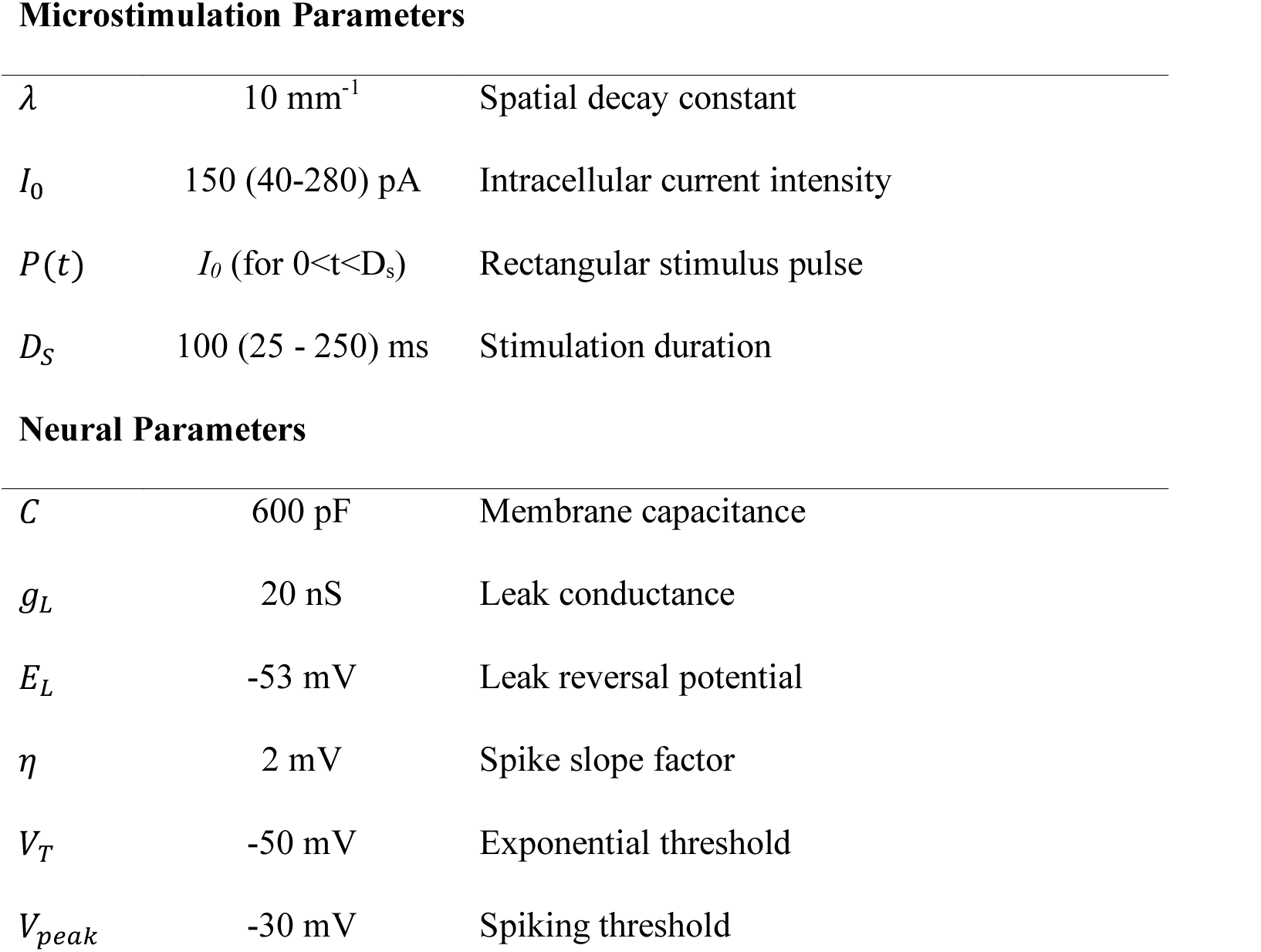
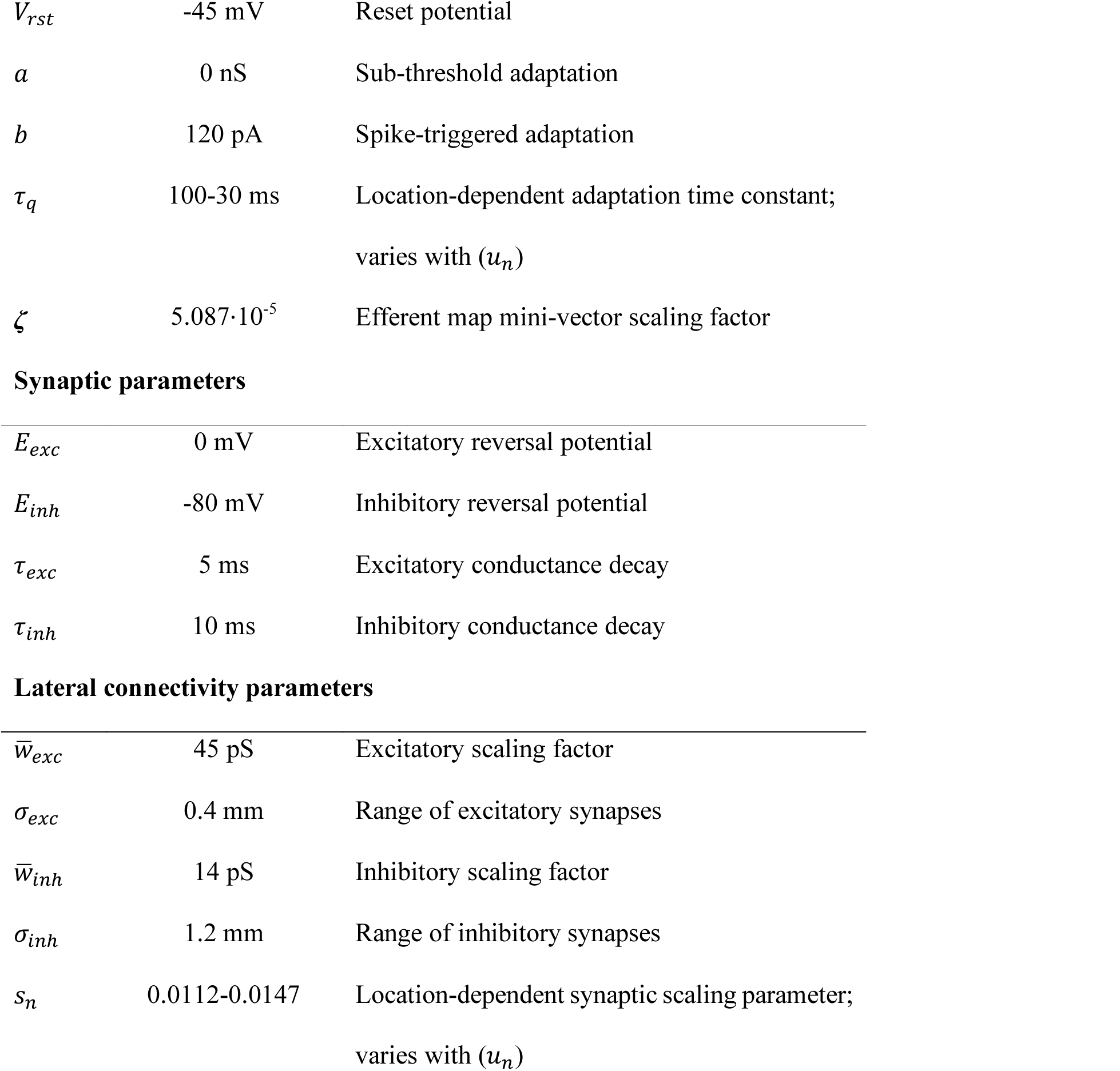
List of all parameters used in the simulations.

Microstimulation Parameters
|  |  |  |
| --- | --- | --- |
| $\lambda$ | 10 mm<sup>-1</sup> | Spatial decay constant |
| $I_0$ | 150 (40-280) pA | Intracellular current intensity |
| $P(t)$ | $I_0$ (for $0 < t < D_s$) | Rectangular stimulus pulse |
| $D_s$ | 100 (25 - 250) ms | Stimulation duration |
| Neural Parameters |  |  |
| $C$ | 600 pF | Membrane capacitance |
| $g_L$ | 20 nS | Leak conductance |
| $E_L$ | -53 mV | Leak reversal potential |
| $\eta$ | 2 mV | Spike slope factor |
| $V_T$ | -50 mV | Exponential threshold |
| $V_{peak}$ | -30 mV | Spiking threshold |
| $V_{rst}$ | -45 mV | Reset potential |
| $a$ | 0 nS | Sub-threshold adaptation |
| $b$ | 120 pA | Spike-triggered adaptation |
| $\tau_q$ | 100-30 ms | Location-dependent adaptation time constant;<br>varies with $(u_n)$ |
| $\zeta$ | $5.087 \cdot 10^{-5}$ | Efferent map mini-vector scaling factor |
| **Synaptic parameters** |  |  |
| $E_{exc}$ | 0 mV | Excitatory reversal potential |
| $E_{inh}$ | -80 mV | Inhibitory reversal potential |
| $\tau_{exc}$ | 5 ms | Excitatory conductance decay |
| $\tau_{inh}$ | 10 ms | Inhibitory conductance decay |

**Table 1** List of all parameters used in the simulations.
|  |  |  |
| --- | --- | --- |
| $\bar{w}_{exc}$ | 45 pS | Excitatory scaling factor |
| $\sigma_{exc}$ | 0.4 mm | Range of excitatory synapses |
| $\bar{w}_{inh}$ | 14 pS | Inhibitory scaling factor |
| $\sigma_{inh}$ | 1.2 mm | Range of inhibitory synapses |
| $s_n$ | 0.0112-0.0147 | Location-dependent synaptic scaling parameter;<br>varies with $(u_n)$ |

The AdEx neuron model employs a smooth spike initiation zone between *V_T_* and *V_peak_*, instead of a strict spiking threshold. Once the membrane potential crosses *V_T_*, the exponential term in Eqn. 7a starts to dominate and the membrane potential can in principle increase without bound. We applied a practical spiking ceiling threshold at *V_peak_* = −30 mV for the time-driven simulations. For each spiking event at time *τ*, the membrane potential is reset to its resting potential, *V_rst_*, and the adaptation current, *q*, is increased by *b* to implement the spike-triggered adaptation:

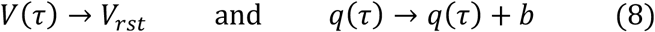

After rescaling the equations, the neuron model has four free parameters (plus the input current) (Touboul, 2008). Two of these parameters characterize the sub-threshold dynamics: the ratio of time constants, *τ_q_*/*τ_m_* (with the membrane time constant *τ_m_* = *C*/*g_L_*) and the ratio of conductances, *a*/*g_L_* (*a* can be interpreted as the stationary adaptation conductance). Furthermore, the resting potential *V_r_* and the spike-triggered adaptation parameter *b* characterize the emerging spiking patterns of the model neurons (regular/irregular spiking, fast/slow spiking, tonic/phasic bursting, etc.).

### 2.3 Current spread function

We applied electrical stimulation by the input current, centered around the site at *[U_E_,****v***_*E*_], according to Eqn. 5. We incorporated an exponential spatial decay of the electric field from the tip of the electrode:

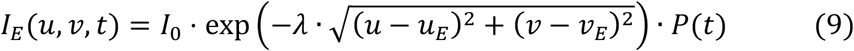

with *λ* (mm^−1^) a spatial decay constant, *I_0_* the current intensity (in pA), and a rectangular stimulation pulse given by *P(t)* = 1 for 0 < *t* < *D*_S_, and 0 elsewhere. Thus, only a small set of neurons around the stimulation site will be directly activated with this input current (see Results). Throughout this paper, we used a fixed input current profile (*I_o_* = 150 pA, *λ*=10 mm^−1^ and D_S_=100 ms) except for the final section, where we explore the effect of microstimulation parameters on the resulting saccade. These parameters were determined by the neural tuning of the AdEx neurons in their bursting regime (see Neural tuning and bursting mechanism section in Results).

#### Remark on the current scale

In SC microstimulation experiments, one typically applies extracellular currents in the micro-Ampère range (10–50 μA) to evoke a saccade. In our simulations, we instead take the effective *intracellularly* applied current, which amounts to only a tiny fraction of the total extracellular current leaving the electrode.

### 2.4 The SC model: synapses and lateral connections

The total input current for an SC neuron, *n*, located at (*u_n_*,***v***_*n*_), is governed by the spiking activity of surrounding neurons, through conductance-based synapses, and by the externally applied electrical stimulation input (Eqn. 9):

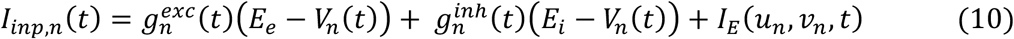

where 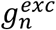 and 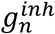 are excitatory and inhibitory synaptic conductances acting upon neuron *n*, *E_e_* and *E_i_* are excitatory and inhibitory reversal potentials respectively. These conductances increase instantaneously for each presynaptic spike by a factor determined by the synaptic strength between neurons, and they decay exponentially otherwise, according to:

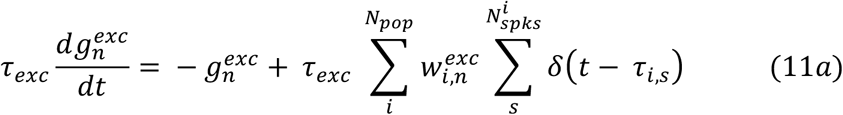

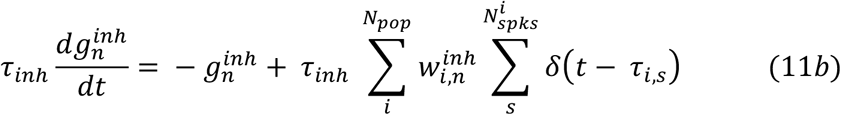

with *τ_exc_* and *τ_inh_*, the excitatory and inhibitory time constants; 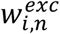 and 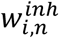 are the intracollicular excitatory and inhibitory lateral connection strengths between neuron *i* and *n*, respectively (Eqn. 12a,b) and *τ_i,s_* is the spike timing of the presynaptic SC neurons that project to neuron *n*. With conductance-based synaptic connections, spike propagation occurs in a biologically realistic way, since the postsynaptic projection of a presynaptic spike depends on the instantaneous membrane potential of the postsynaptic neuron. In this way, the state of a neuron determines its susceptibility to presynaptic spikes.

We incorporated a Mexican hat-type lateral connection scheme in the model, where the net synaptic effect is given by the difference between two Gaussians (Trappenberg, 2001). Accordingly, neurons were connected with strong short-range excitatory and weak long-range inhibitory synapses, which implements a dynamic soft winner-take-all (WTA) mechanism: not only one neuron remains active, but the “winner” affects the temporal activity patterns of the other active neurons. The central neuron governs the population activity, since it is the most active one in the recruited population. As a result, all recruited neurons exhibit similarly-shaped bursting profiles as the central neuron, leading to synchronization of the spike trains within the population (Kasap and Van Opstal, 2017). Two Gaussians describe the excitatory and inhibitory connection strengths between collicular neurons as function of their spatial separation:

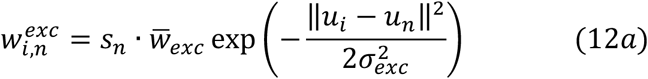

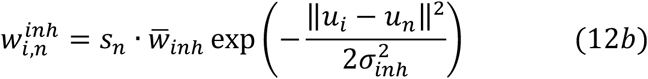

with *w_exc_* > *w_inh_* and *σ_inh_* > *σ_exc_*, and *s_n_* is a location-dependent synaptic weight-scaling parameter, which accounts for the location-dependent change in sensitivity of the neurons due to the variation in adaptation time constants.

### 2.5 Network tuning

Electrophysiological experiments have indicated that the neural responses are well characterized by four principles: (i) a fixed number of spikes for each neuron associated with its preferred saccade vector *N_u,v_* ≅ 20 spikes, (ii) a systematic dependence of the neuron’s cumulative spike count on the saccade vector (dynamic movement field), *N_u,v_*(*R, ϕ, t*), (iii) scaled and synchronized burst profiles of the neurons in the population, resulting in a high cross-correlation, *C_pop_*(*f_n_*(*t*), *f_m_*(*t*)) ≈ *δ_nm_*, between the firing rates of recruited neurons, and (iv) a systematic decrease of the peak firing rate of central neurons in the population, *F_peak_*, along the rostral-caudal axis, together with an increase of burst duration, *T_burst_*, and burst skewness, *S_burst_*.

Goossens and Van Opstal (2012) argued that these properties follow from a systematic tuning of the gaze-motor map, and that they are responsible for the observed saccade kinematics. Here we applied these principles to determine a similarity measure between our simulated responses, and the experimentally recorded gaze motor-map features. In our network model, these features emerge from the interplay between intrinsic biophysical properties of the SC neurons, and the lateral interactions between them.

#### 2.5.1 Distinct biophysical properties

The intrinsic biophysical properties of the neurons were enforced by systematically varying the adaptation time constant, *τ_q,n_*, and the synaptic weight-scaling parameter, *s_n_*, in the motor map. Changes in the adaptive properties of the neurons result in a varying susceptibility to synaptic input. The synaptic weight-scaling parameter corrects for the total input activity. These distinct biophysical properties capture the systematically changing firing properties of SC cells along the rostral-caudal axis of the motor map, while keeping a fixed number of spikes for the neurons’ preferred saccades *N_u,v_*(*R, ϕ*). Following the brute-force algorithm from our recent paper (Kasap and Van Opstal, 2017), the location-dependent [*τ_q,n_*, *s_n_]* value pairs for the neurons were fitted to ensure a fixed number of spikes per neuron under a given microstimulation condition, and the subsequent excitation through lateral interactions (see below, Eqns. 16 and 17). These parameters were first tuned for isolated neurons. The lateral interactions ensured that the bursting profiles in the population remained scaled versions of each other and had their peaks synchronized (evidenced from a high cross-correlation, *C_pop_*, between the burst profiles across the population). The *s_n_* values of Eqn. 12a,b were scaled by the number of neurons in the population.

#### 2.5.2 Lateral connectivity

The single-unit recordings also suggested that for each saccade the recruited population size, and hence its total number of spikes, is invariant across the motor map. The widths of the Mexican-hat connectivity (*σ_exc_* and *σ_inh_*) govern the spatial range of a neuron’s spike influence in the network, and directly affect the size of the neural population. In our model, these widths were fixed, such that they yielded local excitation and global inhibition. The connection strengths (*w_exc_* and *w_inh_*), on the other hand, affect the spiking behavior and local network dynamics, as they control how much excitation and inhibition will be received by each single neuron, and transmitted to others, based on the ongoing activity. Strong excitation would result in an expansion of the population, whereas a strong inhibition would fade out the neural activity altogether. Thus, balanced intra-collicular excitation and inhibition would be required to establish a large, but confined, Gaussian population.

The parameters for the lateral connection strengths were found by a genetic algorithm, as described in our previous paper (Kasap and Van Opstal, 2017). In the current model we used eight saccade amplitudes for each generation to calculate the fitness of each selection (selected as *R=* [2, 3, 5, 8, 13, 21, 33, 55] deg, and *ϕ*=0 deg, to cover equidistant locations on the rostral-to-caudal plane: u= [0.69, 1.08, 1.60, 2.07, 2.56, 3.04, 3.49, 4.00] mm, and v=0 mm, respectively).

The genetic algorithm minimized the root-mean squared errors (RMSE) between the spiking network responses and the rate-based model of Van Opstal and Goossens (2008): from the fitness evaluation for each generation, we calculated the RMSE between the peak firing rates, *F_peak_*; the number of elicited spikes from the central cells in the population, *N_u,v_*(*R, ϕ*); burst durations, *T_burst_*; and burst skewness, *S_burst_*. Furthermore, the cross-correlations, *C_pop_*, between all active neurons and the central cell were included too to ensure that the experimentally observed gaze-motor map characteristics were taken into account for parameter identification. The fitness function was defined by a weighted RMSE summation:

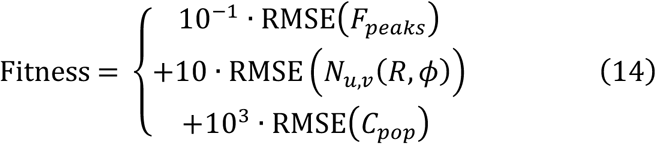

where the weights (0.1, 10, 10^3^) were empirically chosen to cover similar ranges, since the *F_peaks_* vary from roughly 430-750 spikes/s, the number of spikes varies between 18 and 22, and the cross-correlation values are < 1.

Peak firing rates of the central neurons from each population were calculated by convolving the spike trains with a Gaussian kernel (Eqn. 3; 8 ms kernel width), to determine spike-density functions of instantaneous firing rate. RMSE values for *F_peak_* along the rostral-caudal axis of the motor map were subsequently tuned by approximating the following relation:

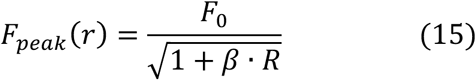

where F_0_ = 800 spikes/s and β = 0.07 ms/deg (taken from Van Opstal and Goossens, 2008). The RMSE of the total spike counts during the burst from the central cells in the population were tuned to *N_u,v_* = 20 spikes, and was required to be independent of the neuron’s position in the map. Synchrony of the neural activity within the recruited population was quantified by the RMSE of deviations for the cross-correlations between the central cell and all other active cells in the recruited population.

### 2.6 Generating eye movements

Eye movements were generated by the population activity following the linear ensemble-coding model of Eqns. 2b and 3. We applied the two-dimensional efferent motor map of Eqn. 5. For any network configuration throughout this paper, the unique scaling factor of the efferent motor map (ζ) was calibrated for a horizontal saccade at (*x,y*) = (21,0) deg. The resulting eye-displacement vector, 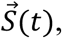 was calculated from the spike trains by interpolation with a first-order spline to obtain equidistant time samples. The interpolated data were further smoothed with a Savitzky-Golay filter, to obtain smooth velocity profiles.

## 3. Results

### 3.1 Neural tuning and bursting mechanism

Figure 1 shows the membrane potential traces for three model neurons, differing in their adaptation time constants, *τ_q_*, which were stimulated under different microstimulation paradigms. The electrical stimulus strength increased from a low amplitude (*I_0_*=50 pA; light blue traces) to a high intensity (*I_0_=250* pA, dark-blue traces), for stimulation durations between 25 and 225 ms. Note that for these different microstimulation regimes, the burst onsets and burst shapes (i.e., the instantaneous firing rates) could differ, even when the number of elicited spikes would be the same. These responses illustrate how the biophysical properties of the neurons affected their bursting behavior.

First, the neuron could respond after the stimulation had terminated. Such a feature, as well as the bursting behavior, is only captured by more complex spiking neuron models. Even when the input current amplitude cannot drive a neuron rapidly to its first spike to initialize the burst (light traces), it suffices if the neuron’s membrane potential crosses a certain threshold (*V_T_* in the AdEx neuron). The neuron can then elicit a spike after the stimulation is over (visible for stimulation durations < 75 ms).

Second, the stimulation amplitude determines the response onset: as the amplitude increases, the first spike occurs earlier. Such a behavior is to be expected, since the neuron model acts as an integrator (Katnani and Gandhi, 2012); higher input currents thus drive a neuron faster to its spiking threshold.

Third, the different neurons respond differently to long stimulation trains (>175 ms). While the neuron with a longer adaptation time constant (*τ_q_* =84.6 ms; Fig.1A) responds with repetitive bursts of 4 to 5 spikes, separated by a silent period, the faster recovering neuron (*τ_q_* =52.4 ms; Fig.1C) elicits more and more spikes after the initial burst, especially for the higher current amplitudes (dark traces).

Interestingly, the neurons with the intermediate (Fig. 1B) and short (Fig. 1C) adaptation time constants switch between different bursting behaviors as the current amplitude increases along with longer stimulation durations. Regular short bursts with silent periods in between result from the slow decay of the adaptation current, which acts on the membrane potential as an inhibitory current. Hence, the adaptation time constant determines how fast a neuron will recover after each spike in a burst. Therefore, the strongly adapting neuron with a long *τ_q_* will require more input current to elicit another spike (Fig. 1A and B for stimulation duration >175 ms), and thus after the fourth spike in the burst, the adaptation current is already high enough to break the bursting cycle. The fast recovering neuron (Fig. 1C, short *τ_q_*) continues its burst with more spikes (dark traces at longer durations (B, C).

A phase plot of the instantaneous adaptation current vs. the membrane potential provides a graphical analysis of the effects of changing the neural parameters, the current input, and the initial state, on the evolution of the dynamical system. Figure 2 shows a number of phase-trajectories for the Adex model, for the parameters used in the simulations of the SC motor map. Nullclines illustrate the boundaries of the vector fields in the AdEx neuron’s phase plane. The *V*-nullcline (*V_null_*; i.e., *dV*/*dt*=*0* for Eqn. 7a) and the *q*-nullcline (*q_null_; i.e., dq*/*dt*=*0* for Eqn. 7b) are shown as gray lines. Fixed points of the system lie at the intersections of these nullclines. A stable fixed point of the system is found at [-53 mV, 0 nA]. In all subfigures that is the starting point of the trajectories, and the state variables of the neurons will converge to this stable fixed point in the absence of input.

**Figure 2:**
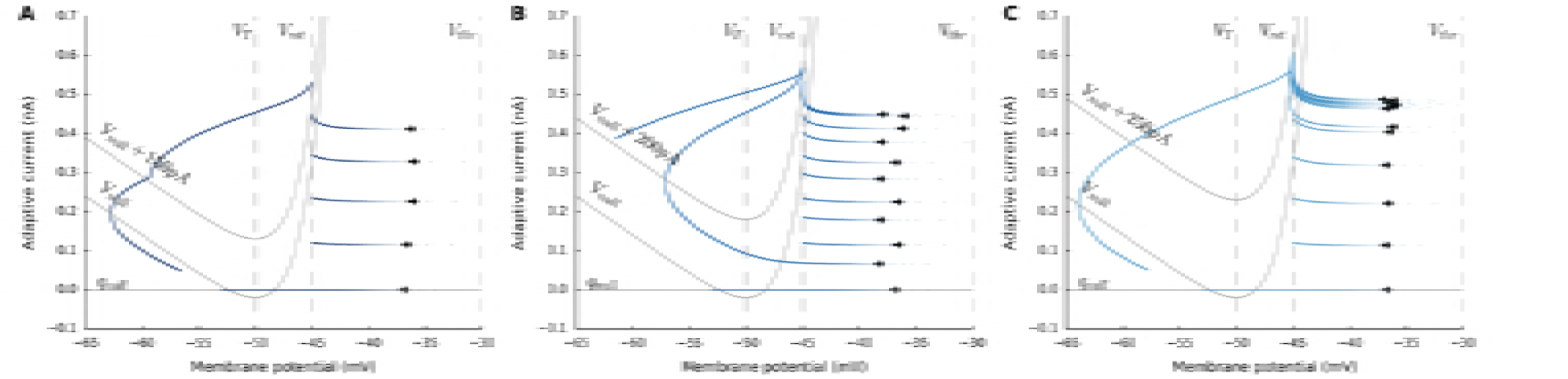
Bursting mechanism of the AdEx neuron model. Phase plots of V(t) vs. q(t) of the neural dynamics of the same three neurons of Figure 1. Biophysical parameters of the neurons were selected for their bursting responses to a ramp stimulus, with varying current amplitude and durations (traces are marked in Figure 1: **A:** a burst with 5 spikes (x); **B:** two burst cycles with 6 and 5 spikes (o); **C:** a burst cycle with more than 10 spikes (+).

The q-nullcline follows a linear trajectory, whereas the V-nullcline represents a convex function because of the superposition of two V-dependent parts. For *V* < *V_T_*, the exponential term can be omitted and the linear *V* dependence will have a slope of *g*_L_. For *V* > *V_T_*, the exponential term will dominate with a sharp increase as *V* increases. When a neuron receives input, the V-nullcline shifts upward by as much as the current density, and the response of the neuron follows a trajectory on the phase plane toward the spiking threshold. The blue trajectories show the evolution of the state variables for three neurons with different *τ_q_* values, and stimulated at different current strengths. The horizontal arrows show the membrane potential in the spike initiation zone, *V* > *V_T_*. Spikes occur when the membrane potential overcomes the spiking threshold, *V* > *V_thr_*. After a spike, the membrane potential is reset, and the adaptation current is increased by *b* (Eqn. 7). The spiking threshold, *V*_thr_, and the reset potential, *V*_rst_, are indicated by the vertical dashed lines. With each spike, the adaptive current increases more and once it reaches values above the *V*-nullcline, the adaptive current is high enough to suppress the neuron from continued bursting, and hyperpolarizes.

In Fig. 2A, the phase trajectory crosses values over *V_null_* + 150 *pA* after 5 spikes. Due to the hyperpolarization, the membrane potential starts to drop. The phase plot shows that the microstimulation is finished when the membrane potential decreases to −58 mV, and the smooth trajectory is seen disrupted. In Fig. 2B, there is a second burst cycle since the microstimulation duration is much longer. After the first burst cycle crosses *V_null_* + 200 *pA* with 6 spikes (arrows are placed closer to *V_thr_*), neuron follows the trajectory to the spike initiation zone for a second burst cycle with 5 spikes. The end of the microstimulation coincides with the second burst cycle and afterwards the membrane potential decreases fast due to the high adaptive current acting on the neuron. In Fig. 2C, the neuron gets stuck in its first cycle and continues spiking repetitively. This pattern is due to the fast decay of the adaptive current, which drops by more than *b* after each spike. Therefore, the neuron would continue spiking repetitively, as long as the current is applied.

The neurons in the network were tuned to respond with a fixed number of spikes in a burst cycle (as in Fig. 2A). This initial burst sets up a large population activity through the lateral connections. *V_null_* fluctuates for each neuron with the network dynamics, depending on the input from other neurons in the population. Microstimulation parameters were chosen such that the central neuron of the population would respond with a burst cycle of 4-5 spikes (typically, *D_S_*=100 ms, and *I_0_* =150 pA), independent of the biophysical properties of the neuron. To that end, the adaptation time constant, *τ_q,n_*, and the synaptic weight-scaling parameter, *s_n_*, for each neuron were determined by applying a fifth order polynomial fit to produce a fixed number of spikes (N=20) for self-exciting neurons:

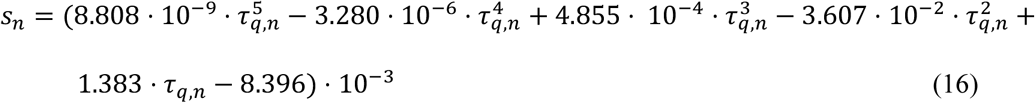

The self-excitation mimics the population activity, since the central cell’s burst profile is representative for the entire population activity, due to burst synchronization across the active neurons. The adaptive time constant, *τ_q,n_*, varied from 100-30 ms in a linear way with the anatomical rostral-caudal location of the neurons, according to:

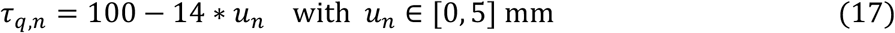

### 3.2 Microstimulation without lateral interactions

The current density drops rapidly with distance from the microelectrode tip, as given by the current spread function (Eqn. 9, with *λ* = 10 mm^−1^, D_S_=100 ms, and *I_0_*=150 pA). Figure 3A illustrates this decay of current density on the motor map surface. The pulsed input current is presented onto the collicular surface at a site corresponding to the visual image point (***u***(***R***), ***v***(***ϕ***) in Eqn. 5; Fig 3B and C). Microstimulation directly activated only a small set of neurons within a 250 ***μ***m radius. Figure 3B and C shows the number of spikes elicited by the activated neurons in the absence of intra-collicular lateral interactions. Each activated neuron elicited only 4-6 spikes within a given input duration range, regardless the electrode’s location. These spikes arose from the initial bursting regime of the neurons until the adaptation current built up with repetitive spikes that canceled the microstimulation input (see Fig. 2). The input amplitude affected the response delay of the neurons between stimulation onset and their first spike. Thus, in the model these small neuronal subsets generated only a brief pulse signal that is supposed to set up the entire population activity through lateral connections.

**Figure 3:**
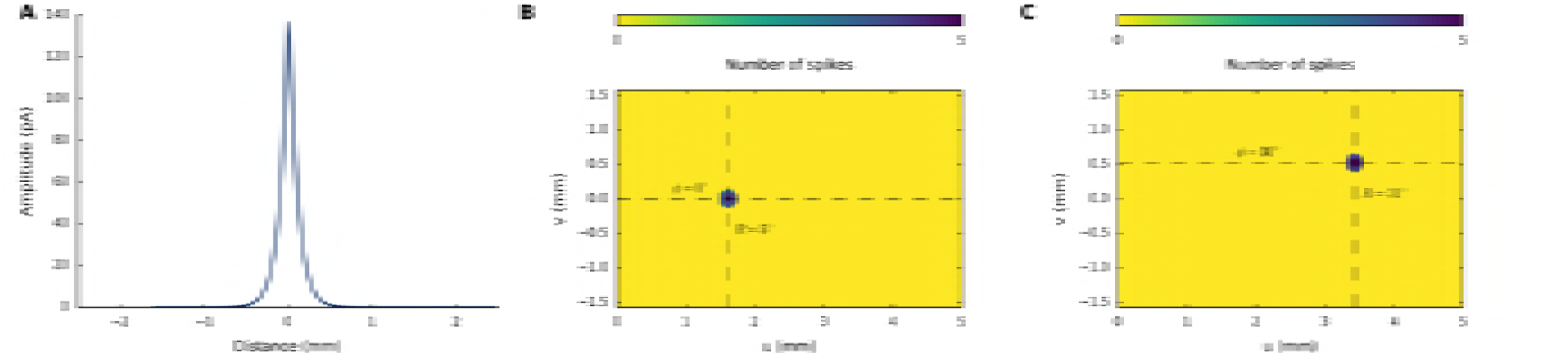
Spatial properties of input current and neural response. **(A)** Input stimulus of 150 pA (100 ms), is presented to the network around the vicinity of the tip of the electrode. Current amplitude drops exponentially away from the tip location at 0 with λ=10 mm^−1^. **(B,C)** Spike counts of neurons activated by microstimulation, without including lateral connections in the motor map. The gaze-motor map is stimulated at the corresponding locations prescribed by the logarithmic afferent mapping function (**B**: R = 5 deg, ϕ = 0 deg; **C**: R = 31 deg, ϕ = 30 deg).

### 3.3 Including lateral interactions

We next tested the collicular network response to the same microstimulation parameters as in Fig. 3, while including the lateral interactions. Figure 4A-C shows the recruited neural population at the rostral stimulation site. Clearly, the number of recruited neurons had increased substantially as a result of the network dynamics. The diameter of the circular population extended to about 1 mm in the motor map. In addition, the cumulative activity elicited by the central cells had now increased from about 5 to 20 spikes. Figure 4B shows the neuronal bursts (top spike patterns) from a number of selected cells in the population, together with the associated spike-density functions. The peak firing rate of the central cells was close to 700 spikes/s and dropped in a regular fashion with distance from the population center. Note also that the cells near the fringes of the population were recruited slightly later than the central cells, but that their peak firing rates were reached nearly simultaneously. Moreover, the bursts all appeared to have the same shape. Figure 4C shows the saccade that was elicited by this neural population, together with its velocity profile. The saccade had an amplitude of 5 deg, reaching a peak velocity of about 200 deg/s.

**Figure 4.**
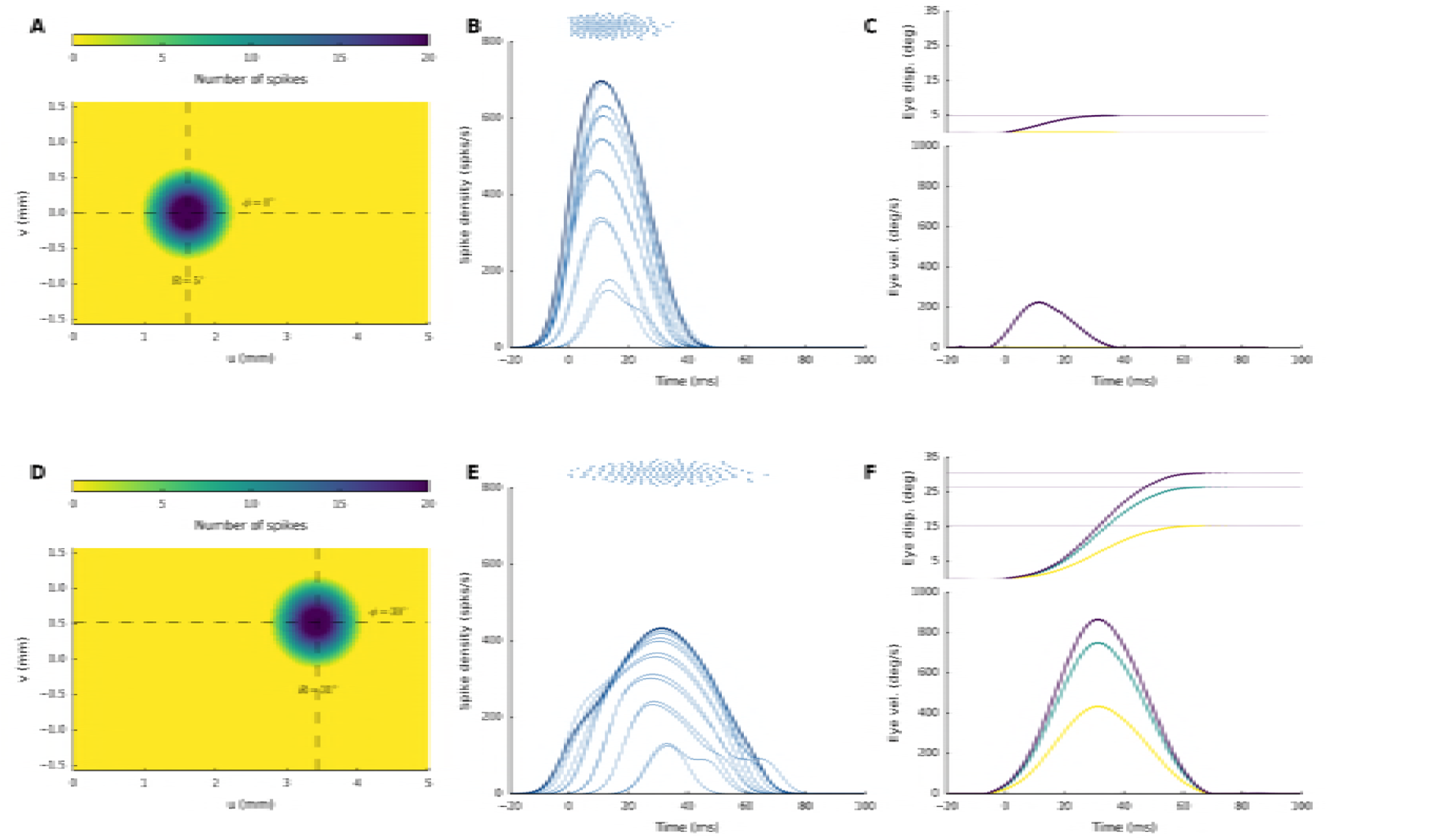
**(A,D)** Spike counts from the gaze-motor map represents the recruited population to microstimulation with lateral interactions. Peak firing rates of the cells decrease with distance from the population center. **(B,E)** Temporal burst profiles of the recruited neurons (taken at 0.1 mm intervals from the central neuron) portray synchronized population activity, here shown along the rostral-caudal direction in the map. Burst durations increase, but the total number of spikes from the population remains the same. **(C,F)** Emerging eye displacements and eye velocity profiles, generated by the linear dynamic ensemble-coding model (Eqns. 2b and 3). Horizontal (green), vertical (yellow), and vectorial (purple) eye-displacement traces.

Figure 4D-F shows the results for stimulation at the more caudal location in the motor map, yielding an oblique saccade with an amplitude of 31 deg. The size of the resulting population activity is very similar to that of the rostral population, and also the number of spikes elicited by the cells is the same. The peak firing rates of the neurons, however, were markedly lower, reaching a maximum of about 450 spikes/s. As a result, the burst durations increased accordingly, from about 50 ms at the rostral site, to more than 70 ms at the caudal site. Note that the saccade reached a much higher peak velocity (about 900 deg/s) than the smaller saccade in Fig. 4C, but its duration was prolonged. Note also that the horizontal and vertical velocity profiles were scaled versions, indicating a straight saccade trajectory.

In Fig. 5 we quantified the collicular bursts in response to microstimulation at different sites along the rostral-caudal axis in the motor map. Figure 5A shows how the evoked collicular bursts of the central cells in the population systematically reduce their peak firing rates, and increase their duration, as the microelectrode moves from rostral (R=2 deg) to caudal sites (R=31 deg). In Fig. 5B we show three major relationships for the bursts of the central cells in the population, for saccade amplitudes between 2 and 65 deg: the peak firing rate (green) drops from about 750 spikes/s to 300 spikes/s, burst duration (purple) increases from about 40 ms to 125 ms, whereas the number of spikes in the burst (light green) remains constant at N=20 spikes. These burst properties, which are due to a precise tuning of the biophysical cell parameters, underlie the kinematic main-sequence properties of saccadic eye movements (Van Opstal and Goossens, 2009; Goossens and Van Opstal, 2012; Kasap and Van Opstal, 2017).

**Figure 5.**
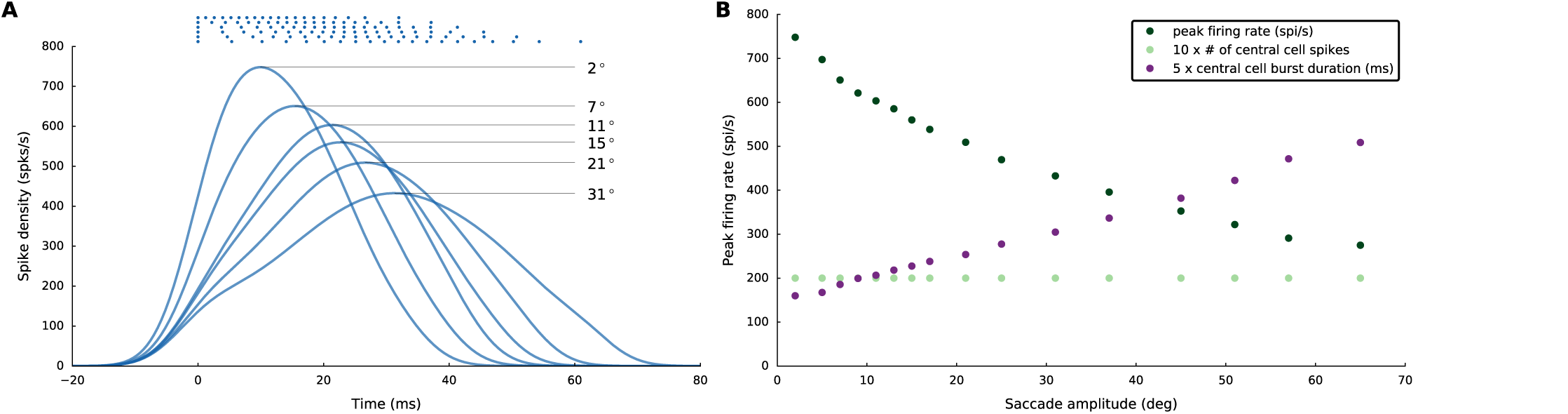
**(A)** Spike trains and burst profiles for the central neurons of different populations (electrode tip positioned at R = 2, 7, 11, 15, 21 and 31 deg). **(B)** Peak firing rates (dark green), number of spikes from the central cells (light green), and the durations of the central cell bursts (purple) for different neural populations between R=2 and 65 deg. Note that the number of spikes for the central cell is constant at about 20 spikes throughout the motor map, while the peak firing rate at caudal sites drops to barely 50% of the rostral stimulation site. Note also that the durations of the central cell bursts increase monotonically with the movement amplitude.

### 3.4 Properties of electrically evoked eye movements

Figure 6A shows the amplitudes and directions of 45 elicited saccades across the 2D oculomotor range (stimulation parameters: *I_0_*=120 pA, *D_S_* = 100 ms). We avoided stimulating near the vertical meridian, as our model included only the left SC motor map (e.g., Van Opstal et al., 1990), and stimulation at very caudal sites (*R*>40 deg), where edge effects of the finite motor map would lead to truncation of the elicited population at the caudal end. Crosses indicate the coordinates of the corresponding motor map locations where stimulation took place; blue dots give the coordinates of the evoked saccade vectors. There is a close correspondence between the motor map coordinates and the elicited saccade vectors. Only for the most caudal sites the saccade vectors tended to show a slight undershoot. We have not attempted to compensate for these minor effects, e.g. by including heuristic changes to the efferent mapping function. The panels of Fig. 6B,C show the evoked saccades for the nine stimulation sites along the horizontal meridian. Note that the saccade duration increased with the saccade amplitude, and that the peak eye velocity showed a less than linear increase with saccade size.

**Figure 6.**
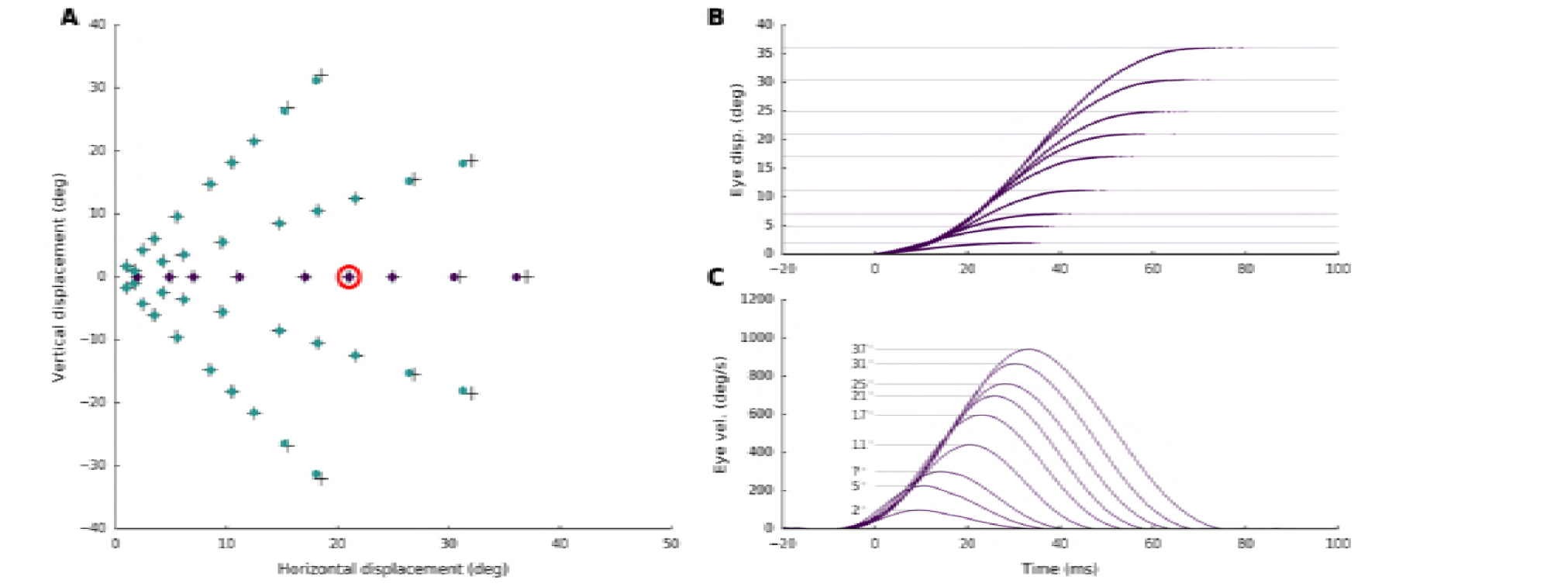
**(A)** Saccade endpoints for stimulation at different sites in the motor map. The scaling parameter of the SC motor map was tuned for a 21 deg horizontal saccade (red circle). **(B)** Eye displacement traces for horizontal saccades (ϕ = deg) [movement amplitudes are highlighted by the thin horizontal lines]. **(C)** Saccadic eye velocity profiles for the corresponding position traces in **B**. Note the clear increase in saccade duration, and the associated saturation of peak eye velocity as function of saccade amplitude.

Figure 7 presents three examples of saccade position and ~ velocity traces for stimulation at sites encoding three different directions, but with a fixed amplitude of *R*=21 deg. The elicited track-velocity profiles are direction-independent. Panels 7B and C also indicate the behavior of the horizontal and vertical saccade components. As these are precisely synchronized with the saccade vector, the ensuing saccade trajectories are straight (not shown).

**Figure 7.**
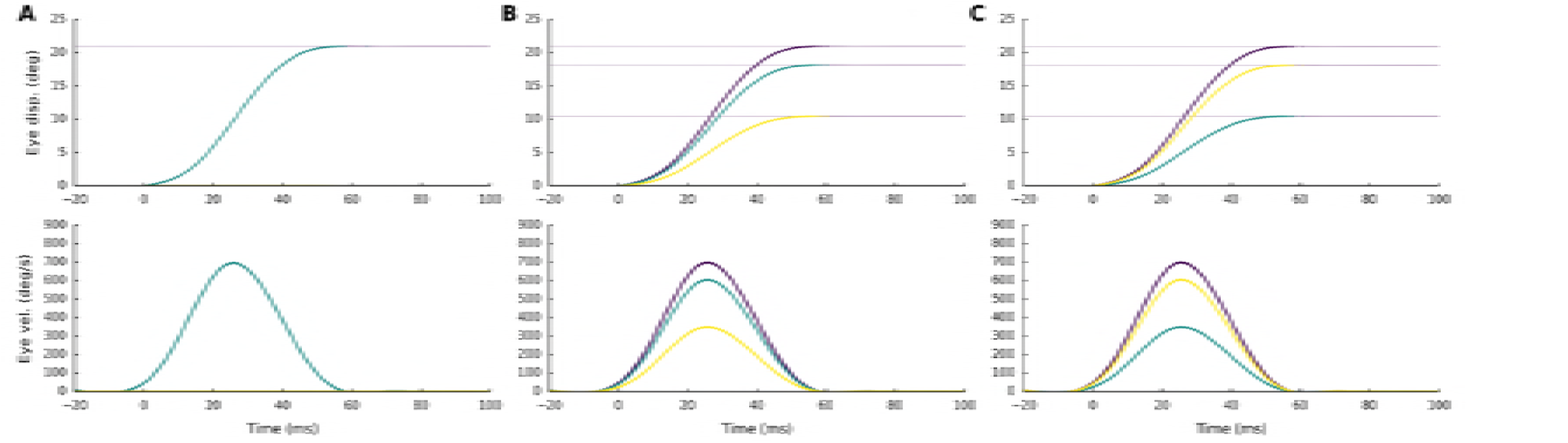
Eye-displacement traces and saccadic eye velocity profiles for three directions (ϕ = 0, 30, 60 deg) **(A, B, C)** with the same amplitude of R = 21 deg. (purple: total vectorial displacement/velocity, green: horizontal, yellow: vertical saccade component).

The main-sequence behavior of the model’s E-saccades is quantified in Fig. 8. Figure 8A shows the nonlinear amplitude vs. peak eye-velocity relationship, described by the following saturating exponential function:

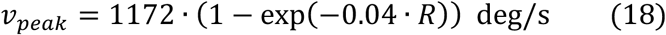

**Figure 8.**
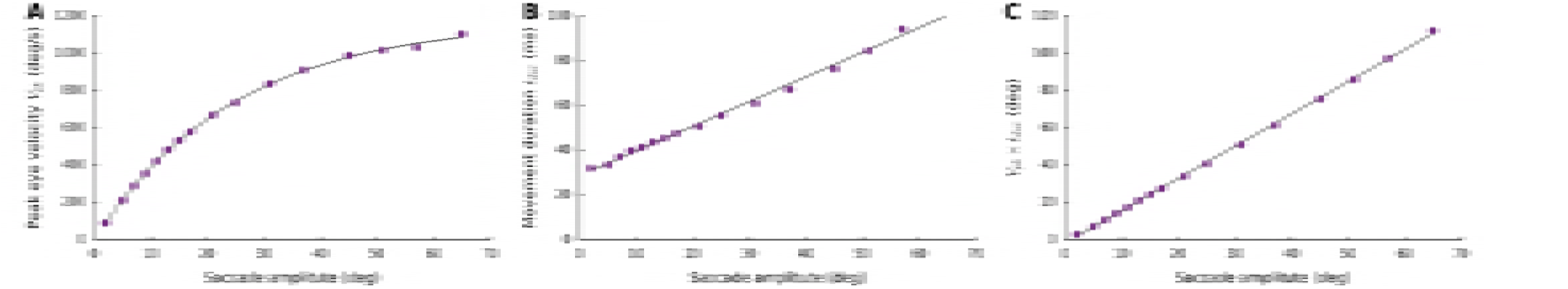
Nonlinear main-sequence behavior of the model, here shown for stimulation at 16 sites along the horizontal meridian of the motor map. **(A)** Saturating amplitude-peak eye velocity relation. **(B)** A straight-line increase of saccade duration with amplitude. **(C)** Saccade amplitude and the product of peak eye velocity and saccade duration, Vpk⋅D, are linearly related with slope, k = 1.7.

From Fig. 8B, the straight-line amplitude-duration relation was approximated to

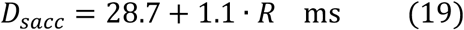

These main-sequence relations were combined into a single, characteristic linear relationship that captures all saccades, normal and slow (Fig. 8C) by:

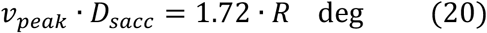

All three relations correspond well to the normal main-sequence properties, as have been reported for monkey and human saccades (e.g., Van Opstal and Van Gisbergen, 1987).

Importantly, the main-sequence behavior of E-saccades was largely insensitive to the applied current strength as soon as it exceeded the stimulation threshold. This feature of the model is illustrated in Fig. 9, which shows E-saccade peak eye-velocity as function of current strength for a fixed stimulation duration of D_S_=100 ms (Fig. 9A). The stimulation was applied at three different sites on the horizontal meridian (corresponding to R=15, 21 and 31 deg). Below *I_0_*=80 pA no movement was elicited, but around the threshold, between 90-120 pA, stimulation evoked slow eye movements, which eventually yielded the final amplitude (Fig. 9B). Immediately above the threshold at 130-140 pA, the evoked movement amplitudes and velocities reached their final, site-specific size (Fig. 9AB), which did not change with current strength over the full range between 140-220 pA. The associated peak eye velocity followed a similar current-dependent behavior for changes in stimulus duration (at a fixed current strength of 150 pA; Fig. 9C). Thus, the quantity that determines evoked saccade initiation is the total amount of current (current amplitude times duration; e.g., Katnani and Gandhi, 2012).

**Figure 9.**
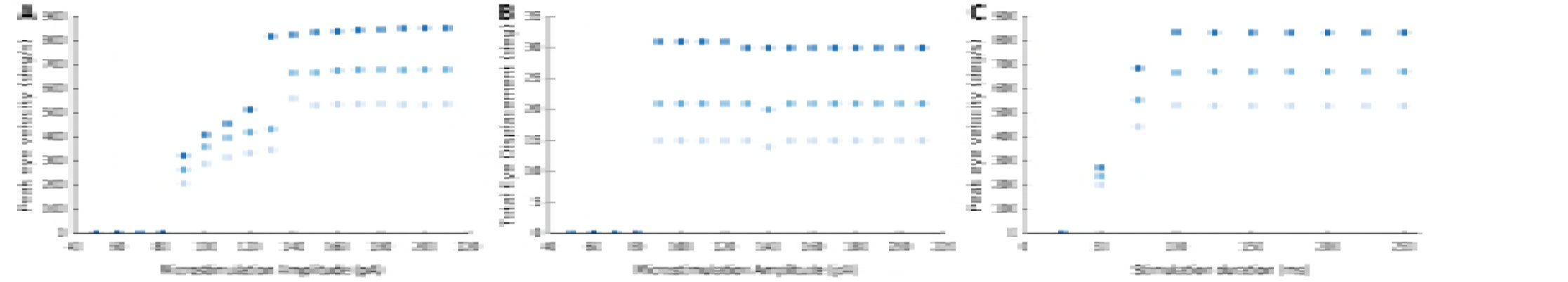
**(A)** Peak eye velocity as function of current strength for stimulation at a site corresponding to R=15 (light), 21 (medium) and 31 (dark) deg, for 100 ms stimulation duration. Beyond the threshold at 140 pA, the evoked eye velocity is virtually independent of the stimulation current. **(B)** Total eye displacement as function of microstimulation strength for stimulation at a site corresponding to R=15 (light), 21 (medium) and 31 (dark) deg for 100 ms stimulation duration. Beyond the threshold at 90 pA, the total eye displacement is independent of the stimulation current. **(C)** Peak eye velocity as a function of microstimulation duration from the same locations at a fixed stimulation strength of 150 pA.

## 4 Discussion

### Summary

The simple linear ensemble-coding model of Eqn. 2b (Van Gisbergen et al., 1987; Goossens and Van Opstal, 2006; Van Opstal and Goossens, 2008) seems inconsistent with the results of microstimulation, when it is assumed that (i) the rectangular stimulation input profile directly dictates the firing patterns of the neural population in the motor map, and (ii) that the neurons are independent, without synaptic interactions.

We here argued that these assumptions are neither supported by experimental observation, nor do they incorporate the possibility that a major factor determining the recruitment of SC neurons is caused by synaptic transmission within the motor map, rather than by direct activation through the electrode’s electric field. We implemented circular-symmetric, Mexican-hat like interactions in a spiking neural network model of the SC motor map and assumed that the current profile from the electrode rapidly decreased with distance from the electrode tip (Fig. 3A). As a consequence, only neurons in the direct vicinity of the electrode were activated by the external electric field (Fig. 3B,C; Histed et al., 2009; 2013).

Once neurons were recruited by the stimulation pulse, however, local excitatory synaptic transmission among nearby cells rapidly spread the activation to create a neural activity pattern which, within 10-15 ms, was dictated by the bursting dynamics of the most active central cells in the population (Fig. 4). As a result, all cells yielded their peak firing rates at the same time, and the burst shapes of the cells within the population were highly correlated. Similar response features have been reported for natural, sensory-evoked saccadic eye movements (Goossens and Van Opstal, 2012), and it was argued this high level of neuronal synchronization ensures an optimally strong input to the brainstem saccadic burst generator to accelerate the eye with the maximally possible innervation.

### Network tuning

The site-dependent tuning of the biophysical parameters of the AdEx neurons, in particular their adaptive time constants and lateral-interaction weightings specified by Eqns. 16-17, caused the peak firing rates of the cells to drop systematically along the rostral-to-caudal axis, while keeping the total number of spikes constant (Fig. 5). As a result, the saccade kinematics followed the nonlinear main-sequence properties that are observed for normal (visually-evoked) saccadic eye movements (Figs. 6-8). In addition, the long-range weak inhibition ensured that the size of the population remained fixed to about 1.0 mm in diameter, and resulted to be largely independent of the applied current strength and the current-pulse duration (Fig. 9).

The lateral excitatory-inhibitory synaptic interactions ensured three important aspects of collicular firing patterns that underlie the saccade trajectories and their kinematics: (i) they set up a large, but limited, population of cells in which the total activity (quantified by the number of spikes elicited by the recruited cells) can be described by a circular-symmetric Gaussian with a width (standard deviation) of approximately 0.5 mm (Fig. 4A,D), (ii) the temporal firing patterns of the central cells (their peak firing rate, burst shape, and burst duration) solely depend on the location in the motor map (Eqn. 15), but the number of evoked spikes remains invariant across the map, and for a wide range of electrical stimulation parameters (Fig. 5), and (iii) already within the first couple of spikes, the recruited neurons all became synchronized throughout the population, in which the most active cells (those in the center) determined the spike-density profiles of all the others (Fig. 4B,E).

Here we described the consequences of this model on the ensuing kinematics and metrics of E-saccades as function of the electrical stimulation parameters. We showed that the network could be tuned such that stimulation at an intensity of 150 pA and a total input current duration of *D_S_* = 100 ms, sets up a large population of activated neurons, in which the firing rates resembled the activity patterns as measured under natural visual stimulation conditions. As a result, the kinematics of the evoked saccades faithfully followed the nonlinear main-sequence relations of normal, visually evoked saccades (Fig. 8). Importantly, above threshold the saccade properties were unaffected by the electrical stimulation parameters (Fig. 9).

### Network normalization

Only close to the stimulation threshold, the evoked activity remained much lower than for supra-threshold stimulation currents, leading to excessively slow eye movements, that started at a longer latency with respect to stimulation onset. Similar results have been demonstrated in microstimulation experiments (e.g. Van Opstal et al., 1990; Katnani and Gandhi, 2012). The saccade peak eye velocity of the model saccades followed a psychometric curve as function of the amount of applied current (Fig. 9). We found that the kinematics of the evoked eye movements at near-threshold microstimulation were much slower than main sequence (Fig. 9). Although this property is readily predicted by the linear summation model (Eqn. 2b), it does not follow from center-of-gravity computational schemes (like Eqn. 2a), in which the activity patterns themselves are immaterial for the evoked saccade kinematics.

Conceptually, the lateral interactions serve to normalize the population activity. Therefore, the total number of spikes emanating from the SC population remains invariant across the motor map, and to a large range of (sensory or electrical) stimulation parameters at any given site. The nonlinear saturation criterion of Eqn. 4 is thus automatically implemented through the intrinsic organization of the SC network dynamics, and do not seem to require an additional downstream ‘spike-counting’ mechanism in order to terminate the saccade response, e.g. during synchronous double stimulation at different collicular sites (see, e.g. Van Opstal and Van Gisbergen, 1989a).

Although other network architectures, relying e.g. on presynaptic inhibition across the dendritic tree, have been proposed to accomplish normalization of the population activity and vector averaging (Van Opstal and Van Gisbergen, 1989a,b; Carandini and Heeger, 1994; Groh, 2001; Van Opstal and Goossens, 2008), substantial anatomical evidence in the oculomotor system to support such nonlinear mechanisms is lacking. We here showed, however, that simple linear summation of the effective synaptic inputs at the cell’s membrane, which is a well-recognized physiological mechanism of basic neuronal functioning, can implement the normalization when it is combined with excitatory-inhibitory communication among the neurons within the same, topographically organized structure. Such a simple mechanism could suffice to ensure (nearly) invariant gaze-motor commands across a wide range of competing neuronal inputs.

### Future work

The two-dimensional extension of our model is a substantial improvement over our earlier one-dimensional spiking neural network model (Van Opstal and Kasap, 2017). It can account for a much wider variety of neurophysiological phenomena. Yet, we have not attempted to mimic every experimental result of microstimulation. A few aspects in our model have not been incorporated yet, or some of its results seem to deviate slightly from experimental observations, which we briefly summarize here.

First, although the network output is invariant across a wide variety of stimulation parameters, and evoked saccade kinematics drop markedly around the threshold (Fig. 9), the present model did not produce small-amplitude, slow movements near the stimulation threshold. This behavior has sometimes been observed for near-threshold stimulation intensities (Van Opstal et al., 1990; Katnani and Gandhi, 2012). In our model, the saccade amplitude behaved as an all-or-nothing phenomenon (Fig. 9B), which is caused by the strong intrinsic mechanisms that keep the number of spikes of the central cells fixed. Although we have not tested different parameter sets at length, we conjecture that a major factor that is lacking in the current model is the presence of intrinsic noise in the parameters and neuronal dynamics that would allow some variability of the evoked responses for small inputs. When near the threshold the elicited number of spikes starts to fluctuate, and becomes less than the cell’s maximum, the evoked saccades will become smaller (and slower) too.

Second, although the main-sequence relations of the model’s E-saccades (Eqns. 18-20) faithfully capture the major kinematic properties of normal eye movements, the shape of the evoked saccade velocity profiles were not as skewed as seen for visually-evoked saccades. As a result, the peak velocity is not reached at a fixed acceleration period, but at a moment that slightly increased with the evoked saccade amplitude (Fig. 6C). We have not attempted to remediate this slight discrepancy, which in part depends on the applied spike-density kernels (here: Gaussian, with width σ=8 ms, Eqn. 3), and in part on the biophysical tuning parameters of the AdEx neurons. However, it should also be noted that a detailed quantification of E-saccade velocity profiles, beyond the regular main-sequence parametrizations (Van Opstal et al., 1990; Katnani and Ghandi, 2012), is not available in the published literature. It is therefore not known to what extent E-saccade velocity profiles and V-saccade velocity profiles are really the same or might slightly differ in particular details.

Third, the electrical stimulation inputs were described by simple rectangular pulses, rather than by a train of short-duration stimulation delta-pulses, in which case also the pulse intervals, pulse durations, pulse heights, and the stimulation frequency would all play a role in the evoked E-saccades (Stanford et al. 1996; Katnani and Ghandi, 2012). We deemed exploring the potential results corresponding to these different current patterns as falling beyond the scope of this study, which merely concentrated on the proof-of-principle that large changes in the input for the proposed architecture of a spiking neural network led to largely invariant results. Note, however, that in our previous paper (Kasap and Van Opstal, 2017) the presumed input from FEF cells to the SC motor map did indeed provide individual spike trains to affect the SC-cells. We there demonstrated that the optimal network parameters could be found with the same genetic algorithm for such spiky input patterns, as applied here (Eqn. 14). The small differences in neuronal tuning parameters for the 1D model with FEF input, compared to the 2D model tuned to electrical pulse input, are mostly due to these fundamentally different input dynamics.

Finally, double-stimulation experiments at different sites within the SC motor map have shown that the resulting saccade vector appears a weighted average between the saccades evoked at the individual sites (Robinson, 1972; Katnani et al., 2012). In the present paper, we have not implemented double stimulation, although an earlier study had indicated that Mexican-hat connectivity profiles in the motor map effectively embed the necessary competition between sites to result in effective weighted averaging (Van Opstal and Van Gisbergen, 1989a). In a follow-up study we will explore the spatial-temporal dynamics of our model to double stimulation at different sites, and at different stimulus onset delays.

## Acknowledgments

This work was supported by the European Commission through FP7 Marie Curie PEOPLE-2012-ITN, project NETT (grant 289146; BK), and by a Horizon 2020 ERC Advanced Grant, project ORIENT (grant 693400; AJvO; BK). The Tesla K40 used for this research was donated by the NVIDIA Corporation.

## References

[1] Bahill AT, Clark MR, Stark L (1975) The main sequence, a tool for studying human eye movements. Math Biosci 204:191–20

[2] Behan M, Kime NM (1996) Intrinsic circuitry in the deep layers of the cat superior colliculus. Vis Neurosci 13: 1031–1042

[3] Brette R, Gerstner W (2005) Adaptive exponential integrate-and-fire model as an effective description of neuronal activity. J Neurophysiol 94: 3637–3642

[4] Carandini M, Heeger DJ (1994) Summation and division by neurons in primate visual cortex. Science 264: 1333–1336

[5] Georgopoulos AP, Schwartz AB, Kettner RE (1986) Neuronal population coding of movement direction. Science 233: 1416–1419

[6] Goossens HHLM, Van Opstal AJ (2012) Optimal control of saccades by spatial-temporal activity patterns in monkey Superior Colliculus. PLoS Comput Biol 8(5): e1002508

[7] Goossens HHLM, Van Opstal AJ (2006) Dynamic ensemble coding of saccades in the monkey Superior Colliculus. J Neurophysiol, 95: 2326–2341

[8] Groh JM (2001) Converting neural signals from place codes to rate codes. Biol Cybern 85: 159–165

[9] Harris CM, Wolpert DM (2006) The main sequence of saccades optimizes speed-accuracy tradeoff. Biol Cybern 95: 21–29

[10] Histed MH, Bonin V, Reid RC (2009) Direct activation of sparse, distributed populations of cortical neurons by electrical microstimulation. Neuron 63: 508– 522

[11] Histed MH, Ni AM, Maunsell JH (2013) Insights into cortical mechanisms of behavior from microstimulation experiments. Prog Neurobiol 103: 115–130

[12] Jürgens R, Becker W, Kornhuber H (1981) Natural and drug-induced variations of velocity and duration of human saccadic eye movements: evidence for a control of the neural pulse generator by local feedback. Biol Cybern 96: 87–96

[13] Kasap B, Van Opstal AJ (2017) A spiking neural network model of the midbrain superior colliculus that generates saccadic motor commands. Biol Cybern 111: 249–268

[14] Kasap B, Van Opstal AJ (2018) Dynamic parallelism for synaptic updating in GPU-accelerated spiking neural network simulations. Neurocomputing, https://doi.org/10.1016/j.neucom.2018.04.007

[15] Katnani HA, Gandhi NJ (2012) The relative impact of microstimulation parameters on movement generation. J Neurophysiol 108: 528–538

[16] Katnani HA, Van Opstal AJ, Gandhi NJ (2012) A test of spatial-temporal decoding mechanisms in the superior colliculus. J Neurophysiology 107: 2442–2452, 2012

[17] Lee C, Rohrer WH, Sparks DL (1988) Population coding of saccadic eye movements by neurons in the superior colliculus. Nature 332: 357–360

[18] Lefèvre P, Quaia C, Optican LM (1998) Distributed model of control of saccades by superior colliculus and cerebellum. Neural Netw 11: 1175–1190

[19] Meredith MA, Ramoa AS (1998) Intrinsic circuitry of the superior colliculus: pharmaco-physiological identification of horizontally oriented inhibitory interneurons. J Neurophysiol 79: 1597–1602

[20] Moschovakis AK, Kitama T, Dalezios Y, Petit J, Brandi AM, Grantyn AA (1998) An anatomical substrate for the spatiotemporal transformation. J Neurosci 18: 10219–10229

[21] Munoz DP, Istvan PJ (1998) Lateral inhibitory interactions in the intermediate layers of the monkey superior colliculus. J Neurophysiol 79:1193–1209

[22] Nickolls J, Buck I, Garland M, Skadron K (2008) Scalable parallel programming with CUDA, AMC Queue 6: 40–53

[23] Olivier E, Porter JD, May PJ (1998) Comparison of the distribution and somato-dendritic morphology of tecto-tectal neurons in the cat and monkey. Vis Neurosci 15: 903–922

[24] Ottes FP, Van Gisbergen JAM, Eggermont JJ (1986) Visuomotor fields of the superior colliculus: a quantitative model. Vision Res 26: 857–873

[25] Phongphanphanee P, Mizuno F, Lee PH, Yanagawa Y, Isa T, Hall WC (2011) A circuit model for saccadic suppression in the superior colliculus. J Neurosci 31: 1949–1954

[26] Phongphanphanee P, Marino RA, Kaneda K, Yanagawa Y, Munoz DP, Isa T (2014) Distinct local circuit properties of the superficial and intermediate layers of the rodent superior colliculus. Eur J Neurosci 40: 2329–2340

[27] Port NL, Wurtz RH (2003) Sequential activity of simultaneously recorded neurons in the superior colliculus during curved saccades. J Neurophysiol 90: 1887–1903

[28] Quaia C, Lefèvre P, Optican LM (1999) Model of the control of saccades by superior colliculus and cerebellum. J Neurophysiol 82: 999–1018

[29] Robinson DA (1972) Eye movements evoked by collicular stimulation in the alert monkey. Vision Res 12: 1795–1808

[30] Robinson DA (1975) Oculomotor control signals. In: Lennerstrand G, Bach-y-Rita P (eds) Basic Mechanisms of Ocular Motility and their Clinical Implications, Pergamon Press, pp 337–374

[31] Scudder A (1988) A new local feedback model of the saccadic burst generator. J Neurophysiol 59: 1455–1475

[32] Smit AC, Van Opstal AJ, Van Gisbergen JAM (1990) Component stretching in fast and slow oblique saccades in the human. Exp Brain Res 81: 325–334

[33] Stanford TR, Freedman EG, Sparks DL (1996) Site and parameters of microstimulation: evidence for independent effects on the properties of saccades evoked from the primate superior colliculus. J Neurophysiol 76: 3360–3381

[34] Tanaka H, Krakauer JW, Qian N (2006) An optimization principle for determining movement duration. J Neurophysiol 95: 3875–86

[35] Touboul J, Brette R (2008) Dynamics and bifurcations of the adaptive exponential integrate-and-fire model. Biol Cybern 99: 319–334

[36] Trappenberg TP, Dorris MC, Munoz DP, Klein RM (2001) A model of saccade initiation based on the competitive integration of exogenous and endogenous signals in the Superior Colliculus. J Cogn Neurosci 13: 256–271

[37] Van Beers RJ (2008) Saccadic eye movements minimize the consequences of motor noise. PLoS One 3(4): e2070

[38] Van Gisbergen JAM, Robinson DA, Gielen S (1981) A quantitative analysis of generation of saccadic eye movements by burst neurons. J Neurophysiol 45: 417–442

[39] Van Gisbergen JAM, Van Opstal AJ, Schoenmakers JJM (1985) Experimental test of two models for the generation of oblique saccades. Exp Brain Res 57: 321–336

[40] Van Gisbergen JAM, Van Opstal AJ, Tax AAM (1987) Collicular ensemble coding of saccades based on vector summation. Neuroscience 21: 541–555

[41] A.J. van Opstal (2016) “The Auditory System and Human Sound Localization Behavior.” Elsevier Publishers, Academic Press, Amsterdam.

[42] Van Opstal AJ, Van Gisbergen JAM (1987) Skewness of saccadic velocity profiles: a unifying parameter for normal and slow saccades. Vision Res 27: 731–745

[43] Van Opstal AJ, Van Gisbergen JAM (1989a) A nonlinear model for collicular spatial interactions underlying the metrical properties of electrically elicited saccades. Biol Cybern 60: 171–183

[44] Van Opstal AJ, Van Gisbergen JAM (1989b) A model for collicular efferent mechanisms underlying the generation of saccades. Brain Behav Evol 33: 90–94

[45] Van Opstal AJ, Goossens HHLM (2008) Linear ensemble coding in midbrain Superior Colliculus specifies the saccade kinematics. Biol Cybern 98: 561–577

[46] Van Opstal AJ, Van Gisbergen JAM, Smit AC (1990) Comparison of saccades evoked by visual stimulation and collicular electrical stimulation in the alert monkey. Exp Brain Res 79: 299–312

[47] Walton MMG, Sparks DL, Gandhi NJ (2005) Simulations of saccade curvature by models that place Superior Colliculus upstream from the local feedback loop. JNeurophysiol 93: 2354–2358

